# Translation of a four-codon ORF in the 3′UTR of *MAP3K10* regulates its expression

**DOI:** 10.64898/2026.04.27.721023

**Authors:** Kirtana Vasu, Swati Ghosh, Sarthak Sahoo, Debraj Manna, Saubhik Som, Md Noor Akhtar, Debaleena Kar, Sandeep M Eswarappa

## Abstract

Several eukaryotic mRNAs are polycistronic because of translatable open reading frames (ORFs) embedded in their untranslated regions (UTRs), in addition to the primary ORF (i.e., the coding sequence). While 5′UTR upstream ORFs (uORFs) are mechanistically and functionally well studied, 3′UTR downstream ORFs (dORFs) remain poorly understood. Here, we identify and characterize a short, translatable dORF in the 3′UTR of *MAP3K10*, which encodes a serine/threonine kinase involved in JNK signalling. A stringent computational screen predicted a conserved RNA G-quadruplex (rG4) within the 3′UTR of *MAP3K10*. Biophysical assays provided more evidence for rG4 formation, which drives translation of a conserved four-codon dORF. Disruption of the rG4 by point mutations or by an rG4-binding ligand reduced dORF translation. Reporter assays using constructs with strategically placed hairpin structures show that translation of the dORF is independent of both the 5’ cap and the translation of the canonical ORF. Notably, deletion of the rG4-dORF module, either in exogenous constructs or in CRISPR-edited cells, led to reduced *MAP3K10* expression. Together, these results provide evidence for the regulation of *MAP3K10* expression by rG4-driven translation of a dORF. Thus, our study contributes to the growing body of evidence suggesting that dORF translation can regulate gene expression.

## INTRODUCTION

Untranslated regions (UTRs) are a key feature of eukaryotic mRNAs. Historically, these regions were considered non-coding segments that do not undergo translation, which led to the term “untranslated regions”. However, accumulating evidence from ribosome profiling and proteomics studies demonstrate that many UTRs are translated and produce short peptides. Many peptides and proteins encoded in the UTRs of several genes have been detected by proteomics studies (1–3). Ribosome profiling studies have revealed footprints of translating ribosomes in the UTRs (4–7). Interestingly, the translation events in the UTRs can have functions independent of the peptides produced (8).

Open reading frames (ORFs) that are translated, termed translons (9), have been identified in both 5′ and 3′UTRs. Among them, translons in the 5′UTRs (also known as upstream ORFs or uORFs) are well-studied. 5′UTR translons regulate gene expression in multiple ways. They can repress the translation of the canonical translon (i.e., the main ORF), regulate translation initiation from alternate initiation sites of the canonical translon, and influence the activity of internal ribosomal entry sites (10–16). Some 5′UTR translons even encode peptides of functional significance (17–20). Regulation of gene expression by 5′UTR translons is important in many physiological and pathological processes, including stress response, circadian rhythm, and carcinogenesis (21–27).

Translons in the 3′UTRs (i.e., translated dORFs) were identified relatively recently (8,28–32). A study based on ribosome profiling predicted that there are 1406 and 1153 3′UTR translons in humans and zebrafish, respectively, more than half of which are initiated from non-AUG codons. Previous studies suggest that translation of dORFs can regulate the expression of their corresponding genes, either positively or negatively. However, the mechanisms that initiate translation within 3′UTR translons remain poorly understood (8,33).

Since the 3′UTR translons are present downstream of their corresponding canonical translons, their translation may require non-canonical mechanisms such as leaky scanning, termination-reinitiation, and cap-independent internal ribosome entry to translate them. In the termination-reinitiation mechanism, the 40S subunit continues to associate with the mRNA after termination due to improper recycling of ribosomal subunits, potentially enabling it to reinitiate scanning and translation at the next available ORF. Leaky scanning from the 5′ cap that misses the initiation site of the canonical ORF may also enable initiation of dORF translation. Such events are reported in the upstream ORFs of multiple eukaryotic mRNAs, including GCN4 in yeast and ATF4 in mammalian cells (34–36). Many viruses use cap-independent translation initiation mechanisms to replicate inside host cells (37). Several stress-responsive eukaryotic mRNAs use cap-independent translation mechanisms during various cellular stresses. *Cis*-acting RNA elements known as internal ribosome entry sites (IRESs) represent one mechanism of cap-independent translation initiation (38–40). IRESs are made up of complex RNA secondary structures that directly or indirectly recruit ribosomal subunits for translation initiation.

RNA G-quadruplexes (rG4s) are one type of structure that can mediate cap-independent translation (41). rG4s are formed in guanine-rich sequences when four guanines form planar structures called G-tetrads, which stack on top of each other stabilized by monovalent cations such as potassium. In addition to promoting cap-independent translation initiation, rG4s have been shown to regulate alternative splicing, polyadenylation, mRNA localization, translation elongation, telomere length, biogenesis, and function of miRNAs and piRNAs (42). In this study, we show that the 3′UTR of mammalian *MAP3K10* (aka *MLK2*) mRNA, which encodes a serine/threonine kinase involved in JNK signalling, has an rG4 structure, which drives the translation of a four-codon-long dORF. Our experiments using cells lacking the dORF showed that translation from this dORF regulates the levels of *MAP3K10* mRNA.

## MATERIALS AND METHODS

### Bioinformatics screening for transcripts containing rG4 in the 3′UTR

The human transcriptome was retrieved from NCBI Genome Reference Consortium (GRCh38.p12, accession # GCF_000001405.38). Predicted mRNAs were not included in the analysis. Out of them, 16,628 mRNAs had at least one stop codon in the 3′UTR in-frame with the canonical stop codon. Of these, only the subset of mRNAs having at least one ‘GG’ motif within the region between the two stop codons (canonical and the downstream) were considered. The first and last ‘GG’ motif within this region were determined for all mRNAs. The sequences encompassing 100 nucleotides upstream of the first GG and up to 100 nucleotides downstream of the last GG served as input sequences for the G4-mining tools.

Sequences were first passed through the ‘QuadBase2’ prediction tool (43). Parameters were set as follows: Medium stringency (G3, L1-7), non-greedy algorithm type, plus-strand search with a bulge size of 2. The non-overlapping sequences that gave positive results in QuadBase2 were passed through the ‘G4RNA Screener’ prediction tool (44) with default search parameters (Window size 60, step size 10) and threshold parameters (cGcC score > 4.5, G4H score > 0.9, G4NN > 0.5). Sequences were sorted in Excel by descending order of the G4 scores in the given priority of G4NN>G4H>cGcC. To increase the stringency of screening, sequences with G4NN score less than 0.9 were removed. Duplicate entries from a single accession number were removed, leaving us with 232 candidates. This list of potential G4-forming candidates was compared with the list of genes that showed evolutionary conservation at the 3′UTR (performed previously to predict potential stop codon readthrough candidates (45)). This yielded 11 candidate mRNAs with a predicted G-quadruplex structure and translation in the 3′UTR.

### Nucleotide sequence alignment

The sequences corresponding to the 3′UTR (including the canonical stop codon) of *MAP3K10* from various mammals were retrieved from NCBI and aligned using the Clustal Omega software. The NCBI IDs of the sequences shown in the alignment are *Homo sapiens* (NM_002446.4), *Pan troglodytes* (XM_016935958.4), *Bos taurus* (XM_024979088.2), *Canis lupus familiaris* (XM_038656969.1), *Panthera pardus* (XM_019432140.2), *Felis catus* (XM_023245041.2), *Sus scrofa* (XM_021094377.1) and *Ursus arctos* (XM_026482306.4).

### Construction of plasmids

*Constructs used for in vitro transcription to investigate rG4 structures:* Sequences corresponding to the 3′UTR of *MAP3K10* (1-60 nucleotides or 183-242 nucleotides downstream of the stop codon) were cloned between *BamH*I and *Xho*I sites of pcDNA 3.1/myc-HisB vector. The TERRA sequence that consisted of 10 repeats of the ‘TTAGGG’ motif was cloned in the same enzyme sites.

*Constructs used to investigate the translatability of dORF:* All constructs were made in pcDNA 3.1/myc-HisB vector. *MAP3K10* cDNA was amplified from HEK293 cells. ATG followed by the partial CDS of *MAP3K10* (717 nucleotides of the 3′ end), its canonical stop codon followed by 261 nucleotides of its 3′UTR, were cloned between *Hind*III and *BamH*I sites upstream of and in-frame with the CDS of firefly luciferase (FLuc), which was cloned between *BamH*I and Not*I* sites. The downstream in-frame stop codon in the 3′UTR of *MAP3K10* and the start codon of FLuc were not included in the construct. A linker sequence (5′ GGCGGCTCCGGCGGCTCCCTCGTGCTCGAG 3′) was included between *MAP3K10* and FLuc sequence. For the fluorescence and western blotting-based assays, ATG followed by FLAG tag was cloned between *Hind*III and *Kpn*I sites. The partial coding sequence (717 nucleotides) of *MAP3K10*, its canonical stop codon, and 261 nucleotides of the 3′UTR were cloned between *Kpn*I and *BamH*I sites. The linker sequence mentioned above, followed by the CDS of green fluorescent protein (GFP) without a start or stop codon, was cloned between the *BamH*I and *EcoR*I sites. The HA tag (2x) followed by a stop codon was inserted between the *EcoR*I and *Xho*I sites. The FLAG tag, *MAP3K10*, GFP, and HA tag were all in the same frame.

*Constructs used to investigate the translation initiation activity*: The CDS of *Renilla* luciferase was cloned between the *Hind*III and *BamH*I sites. The coding sequence of firefly luciferase, excluding the start codon, along with the linker sequence, was cloned between the *Xho*I and *Apa*I sites. The 3′UTR rG4 sequence was cloned between *BamH*I and *Xho*I sites. As a negative control, a non-specific sequence (5′ GCGGTGCACAATCTTCTCGCGCAACGCGTCAGTGGGCTGATCATTAACTATCCGCTGGATGACCAGGATGCCATTGCTGTGGAAGCTGCCTGCACTAATATG 3′) was cloned in place of the 3′UTR. The sequence of G4 mutant^G>A^ insert was (5′ GCCCACCCTGCACTG**A**G**A**G**A**A**A**G**A**TGG**A**CAG**A**GATACTCAG**A**GACAG**A**GCAT CATGG**AA**GATTTGGCACAAAATGGAGCAT 3′).

*Constructs used to test for leaky scanning/ reinitiation:* The 42-nucleotide-long hairpin-forming sequence (5′ GCGGTCCACCACGGCCGATATCACGGCCGTGGTGGACCG CAA 3′)(46) or a length-matched control consisting of ‘CAA’ repeats was cloned into the vector backbone. The 5′ hairpin or 5′ control sequences were cloned between the *Hind*III sites upstream of the start codon of RLuc or *MAP3K10* as shown in Figure 5. The 3′ hairpin or 3′ control sequence, followed by the 3′UTR of *MAP3K10* (includes rG4 and dORF), was cloned into the dual luciferase-based vector backbone between *BamH*I and *Xho*I sites. The entire fragment was cloned downstream of the stop codon of RLuc.

*Constructs used to assess the effect of dORF translation*: Partial coding sequence (717 nucleotides) of *MAP3K10*, its canonical stop codon, and 264 nucleotides of the 3′UTR were cloned between *Not*I and *EcoR*I sites of pIRESneo-FLAG/HA vector.

All mutations were generated by PCR-based site-directed mutagenesis and confirmed by Sanger sequencing.

### *In vitro* transcription

The template DNA was linearized using *Xho*I enzyme and subjected to *in vitro* transcription using T7 RNA Polymerase (Thermo Scientific) overnight at 37°C. Samples were treated with RNase-free DNase I (Thermo Scientific) for 2 h at 37°C to remove the template DNA. The RNA was then purified using the miRNeasy Kit (Qiagen) and quantified using Biophotometer (Eppendorf). To confirm the integrity of the RNA, ~ 200 ng RNA was incubated with formamide-containing loading dye and run on a native agarose gel along with RiboRuler High Range RNA Ladder (Thermo Scientific).

### Circular dichroism

RNA was resuspended in Tris-EDTA-containing buffer (10 mM Tris, 1 mM EDTA) either in the presence or absence of KCl (100 mM) to achieve a final concentration of 5 μM. The RNA was denatured at 95°C for 5 minutes and cooled on ice overnight to enable folding. CD spectrum was recorded using the Jasco-J810 spectropolarimeter equipped with Peltier Temperature Controller. The spectrum was recorded between 200-320 nm using quartz cuvette of length 1 mm (Lab Art Optics). Parameters used were as follows: 100 mdeg sensitivity, 1 nm data pitch, 100 or 200 nm/min scanning speed, 2 or 4 sec response time with a band width of 2 nm. Data shown is an average of three accumulations from which spectral contribution of the corresponding buffer has been subtracted.

For the CD melting experiments, the change in ellipticity at 264 nm was measured as a function of temperature (20°C to 95°C). A temperature slope of 2°C/minute was utilised with data points being recorded after every 1°C. The buffer spectrum was subtracted from the sample spectrum, and the resulting curve was subjected to adaptive smoothing (convolution width 5 and deviation noise of 25). This was then normalized using the formula (Ellipticity(t) – min)/(max – min), in which ellipticity(t) is the ellipticity at a temperature ‘t’, max is the maximum ellipticity value at 264 nm, and min is the minimum ellipticity value at 264 nm in the temperature range recorded.

### Mobility shift assay

The *in vitro* transcribed RNA (4 μg) was resuspended in TE buffer. Samples were heated at 95°C for 5 minutes and kept on ice overnight. The RNA marker, RiboRuler High Range RNA Ladder (Thermo Scientific), was heated at 70°C for 10 minutes and then kept on ice for 3 minutes prior to loading. Samples were mixed with non-denaturing loading dye and subjected to 7% polyacrylamide gel electrophoresis in Tris-borate EDTA (TBE) buffer at 110 V for 2.5 h on ice. Gels were stained with ethidium bromide and then visualised using Biorad ChemiDoc Imaging System.

### Cell culture

HEK293 cells were cultured in Dulbecco’s Modified Eagle’s Medium (HiMedia) supplemented with 10% fetal bovine serum (FBS, Gibco) and 1% antibiotics (10000 U/mL penicillin, 10 mg/mL streptomycin, Sigma). Cells were maintained at 37°C in a humidified atmosphere with 5% CO2. Cell identity was confirmed by STR profiling. Cells were tested for mycoplasma contamination twice a year.

### Luminescence-based translation assays

HEK293 cells were transfected with 500 to 750 ng of indicated constructs expressing firefly luciferase and/or *Renilla* luciferase (25-50 ng of pRL-SV40, when transfected separately) at 80-90% confluency in 24-well plates using Lipofectamine 2000 (Invitrogen). Cells were lysed after 24 h using passive lysis buffer. Luciferase activity was measured using the Dual Luciferase Reporter Assay System (Promega) in the GloMax Explorer (Promega). The relative luciferase activity was calculated as the ratio of the activity of firefly luciferase to that of *Renilla* luciferase.

### Western blotting

HEK293 cells at 70-80% confluency were transfected with 4-8 μg of the indicated constructs in 6-well plates using Lipofectamine 2000 (Invitrogen). Cells were harvested 24 h or 72 h after transfection and lysed in lysis buffer containing 20 mM Tris-HCl, 150 mM NaCl, 1 mM EDTA, 1mM EGTA, 1% Triton-X with protease inhibitor cocktail (Promega). Lysates were quantified using Protein Assay Dye Reagent (BioRad) and lysates containing 30 to 50 μg of total protein were boiled in Laemmli buffer at 95°C, then loaded on 10% denaturing SDS-PAGE.

Proteins were transferred onto a 0.45 μ PVDF membrane (Merck) using a Trans-Blot Semi-Dry Electrophoretic Transfer Cell (BioRad). Blocking was performed with 5% skim milk in phosphate-buffered saline containing 0.1% Tween 20. The membrane was incubated overnight with the primary antibody (Anti-HA antibody Roche # 11867423001 and anti-Actin antibody Sigma # A3854) diluted in blocking buffer or PBS. Following washes, the membrane was probed with peroxidase-conjugated secondary antibody (Jackson ImmunoResearch # 712-035-153). Blots were developed with Clarity ECL reagent (Bio-Rad) or Femto Glow HRP Substrate (Giri Diagnostics), and images were captured with a ChemiDoc Imaging System (Bio-Rad). Band intensities were quantified using ImageJ.

### Flow cytometry

HEK293F cells were seeded in 24-well plates to achieve 70–80% confluency at the time of transfection. Cells were transfected with 2.5 μg of constructs carrying an N-terminal FLAG tag and a C-terminal GFP-HA tag. Transfections for each construct were done in triplicate using Lipofectamine 2000 (Invitrogen) in serum-free, antibiotic-free OptiMEM (Gibco). Cells were subjected to flow cytometry at 72 h post-transfection using Cytoflex LX (Beckman Coulter). A total of 50,000 events were measured per sample. Analysis was done using CytExpert Software.

### RNA isolation and RT-PCR

Total RNA was isolated using RNAiso Plus reagent (TaKaRa) according to the manufacturer’s instructions. RNA quality and concentration were measured using Biophotometer (Eppendorf), and an equal amount of RNA (1 - 2.5 μg) was used for cDNA synthesis. First-strand cDNA synthesis was carried out using Oligo(dT) primers and RevertAid Reverse Transcriptase enzyme (Thermo Scientific) according to the manufacturer’s instructions.

Primer sequences (5′-3′) used for PCR are given below:

FLuc: CAACTGCATAAGGCTATGAAGAGA & ATTTGTATTCAGCCCATATCGTTT

*ACTB*: CACCAACTGGGACGACAT & ACAGCCTGGATAGCAACG

GFP: AAGTTCATCTGCACCACCG & TCCTTGAAGAAGATGGTGCG

### Quantitative real-time PCR

Total RNA was isolated as described above. 1.5 μg RNA was used for first strand cDNA synthesis using Oligo(dT) primers as described above. TB Green Premix Ex TaqII (TaKaRa) was used to perform qRT PCR in the CFX96 real-time PCR system (Bio-Rad). For amplification of endogenous *MAP3K10* and *ACTB*, shuttle PCR protocol was used: 95°C for 5 min, 40 cycles of (95°C for 30 s and 64°C for 30 s), followed by final extension at 64°C for 5 min. For amplification of cDNA from cells transfected with N-terminal FLAG HA-tagged *MAP3K10* constructs, the following PCR conditions were used: 95°C for 5 min, 40 cycles of (95°C for 30 s, 55°C for 30 s, and 72°C for 30 s), followed by a single final extension step at 72°C for 5 min. Melt curves were generated after the reaction and the amplification products were visualised by agarose gel electrophoresis to make sure that a single amplicon of expected size is obtained. Relative expression of the target mRNA relative to that of *ACTB* was calculated using the 2^-ΔΔCt^ method. Primer sequences (5′ to 3′):

*MAP3K10*: AAGTGGGACATGGAGCCGCG & CCAGGTTGGGGGCACTTGATGAC

FLAG-HA-tagged *MAP3K10*:

TAATACGACTCACTATAGG & ATGCGCGGCCGCTTAAGCGTAATCGGGCAC

*ACTB*: CACCAACTGGGACGACAT & ACAGCCTGGATAGCAACG

### *In vitro* transcription and translation

5 μg of firefly luciferase-encoding reporter plasmids were linearised using *Not*I enzyme. After purification, 1 μg of linearised plasmid was subjected to *in vitro* transcription using T7 RNA Polymerase (Thermo Scientific) at 37°C for 3 h. RNA was purified using GeneJET RNA purification kit (Thermo Scientific) and quantified using BioPhotometer (Eppendorf). Agarose gel electrophoresis in TAE buffer in the presence of formamide-containing dye was used to confirm the integrity of the transcribed RNA. 1-2 μg of purified RNA was *in vitro* translated using rabbit reticulocyte lysate (Promega) at 30°C for 2 h as per the manufacturer’s instructions. Luciferase activity was then measured as described above.

### Generation of cells with deletions in the 3′UTR of *MAP3K10* using CRISPR-Cas system

The sgRNAs (5′ GATGGGGGATTTGGCACAAAA 3′ and 5′ GGCACAAAATGGAGCAT TAA 3′) were cloned in the pSpCas9(BB)-2A-GFP (PX458) plasmid. 2 μg of sgRNA-plasmids were transfected into HEK293 cells at 75% confluency in 6-well plates using Lipofectamine 2000. GFP-positive single cells were sorted into 96-well plates 48 h after transfection using FACSAria™ II sorter (BD Biosciences). The clones were expanded and then screened for the presence of deletions by genomic DNA PCR using primers flanking the expected deletion site (5′ GGCCTGCCCACCACCGCCCG 3′ and 5′ GGCAACAGAGCAAGACTCTGTCTCC 3′). The amplicons were sequenced to identify the exact deletions or mutations.

### Statistics

Unpaired Student’s t-test was used to find out the statistical significance. Welch’s correction was applied whenever the variance was different across datasets. Paired Student’s t-test was used for the quantitative real-time PCR assays and the densitometry analysis of western blot images. Information on the experimental replicates and P values are provided in the figure legends.

## RESULTS and DISCUSSION

### The 3′UTR of *MAP3K10* has a predicted RNA G quadruplex (rG4) structure and an evolutionarily conserved downstream open reading frame (dORF)

We set out to identify human mRNAs that exhibit 3′UTR translation regulated by rG4 structures. We employed a bioinformatic workflow outlined in Figure 1A. The 3′UTR can be translated either by stop codon readthrough (SCR) or by translation of a dORF (8,47). As the stop codon is a common feature in both cases, we shortlisted all human mRNAs with 3′UTR stop codons that are in frame with their respective canonical stop codons. This was followed by the identification of a potential rG4 structure in the 3′UTR using two computational tools - QuadBase2 and G4RNA Screener (43,48). This screening identified 232 mRNAs harbouring high-confidence rG4 structures (G4 Neural Network Score, G4NN score > 0.9) in their 3′UTR. This list of mRNAs was compared with the list of mRNAs exhibiting evolutionary conservation in the 3′UTR at the predicted amino acid level, which was performed previously (45) (Fig. 1A). This comparison revealed 11 distinct mRNAs, including *MAP3K10* (also known as *MLK2*) (Table S1). *MAP3K10* encodes a Mitogen-Activated Protein (MAP) Kinase Kinase Kinase, a member of the serine/threonine kinase family.

**Figure 1.**
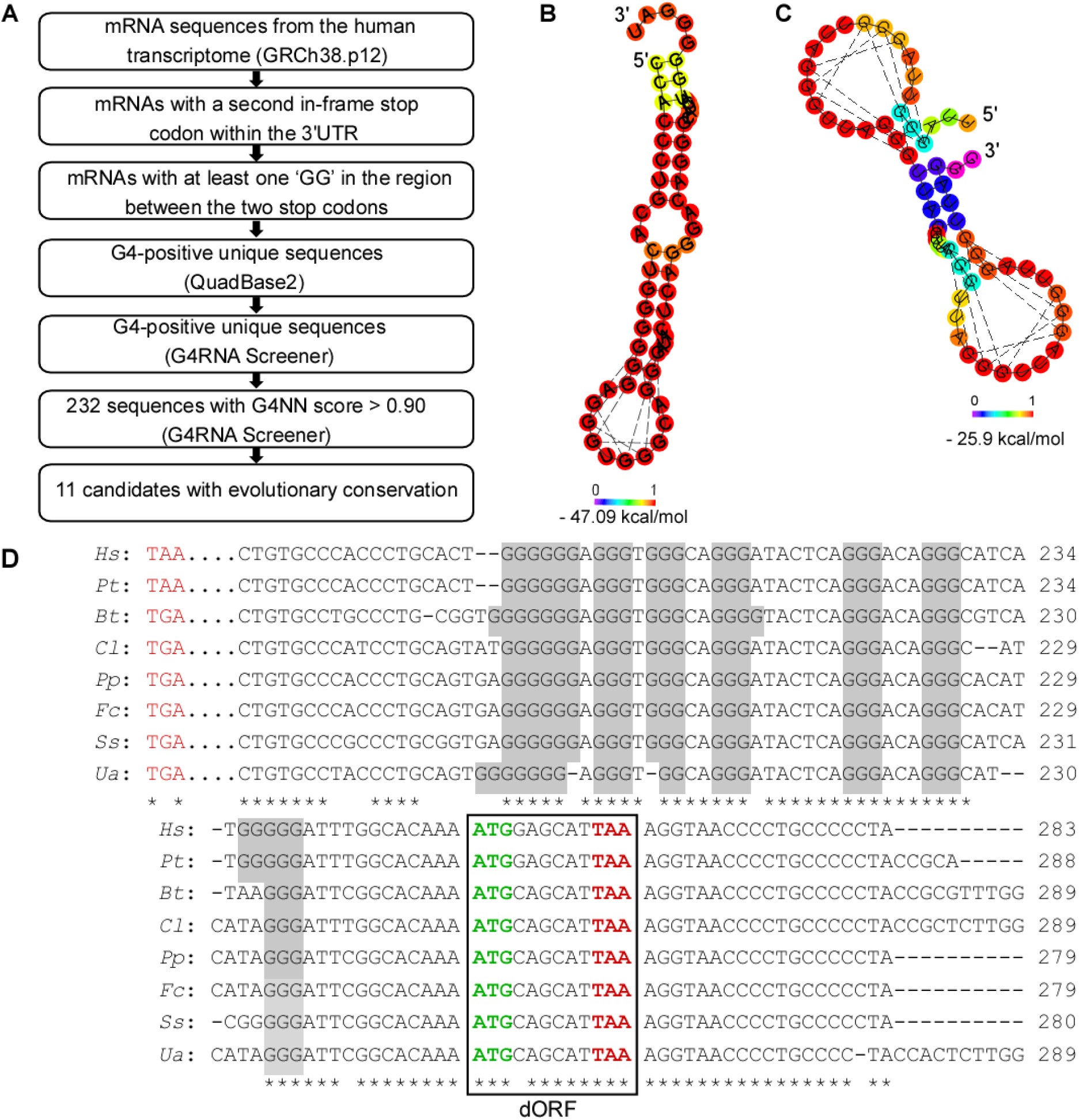
The 3′UTR of *MAP3K10* contains a putative RNA G-quadruplex (rG4) and a conserved downstream ORF (dORF). (A) Flowchart showing the steps in the bioinformatics screen used to identify candidate mRNAs having putative G4-forming sequences in their 3′UTR. (B and C) rG4 structure predicted in the 3′UTR (183 to 242 nucleotides) of *MAP3K10* (B) and TERRA (C). RNAfold web server was used for the prediction. Dotted lines indicate Hoogsteen base pair interactions in the predicted rG4. Color code indicates the probability of the base pairing. (D) Alignment of distal nucleotide sequences of the 3′UTR of *MAP3K10* from multiple mammalian species. The numbers indicate the position of the last nucleotide in a line from the canonical stop codon (position 1 being immediately after the canonical stop codon). Seven G-stretches are highlighted. The dORF with start and stop codons is highlighted within a box. * indicates 100% conservation within the sequences shown. Hs, *Homo sapiens*; Pt, *Pan troglodytes*; Bt, *Bos taurus*; Cl, *Canis lupus familiaris*; Pp, *Panthera pardus*; Fc, *Felis catus*; Ss, *Sus scrofa*; Ua, *Ursus arctos*.

RNAfold webserver also predicted rG4 in the nucleotide sequence from 183 to 242 (G4NN score 0.97) in the 3′UTR of *MAP3K10* (49). The estimated free energy of this structure was −47.09 kcal/mol (Fig. 1B). Telomeric repeat-containing RNA (TERRA, 10 repeats of ‘UUAGGG’), which has a known rG4 structure, served as a positive control. It also showed a predicted rG4 with an estimated free energy of −25.9 kcal/mol (Fig. 1C) (50). Alignment of *MAP3K10* 3′UTR sequences from multiple mammalian species revealed seven evolutionarily conserved stretches of Gs (≥ 3 Gs), which is consistent with a potential rG4 structure (Fig. 1D and S1A). Notably, the same region in the 3′UTR of *MAP3K10* was identified as a potential rG4-forming sequence in computational as well as experimental screens conducted by other groups, which provides independent support to our predictions (51,52).

In addition to the rG4 sequence, the multiple sequence alignment revealed an evolutionarily conserved short dORF of just four codons (including start and stop codons) in the *MAP3K10* 3′UTR. This dORF is in frame with the canonical CDS (i.e., the coding sequence of *MAP3K10*) (Fig. 1D and S1A). Based on these observations, we hypothesized that the 3′UTR of *MAP3K10* is translated, and that the predicted rG4 structure may play a role in this process. We tested these hypotheses experimentally.

### Experimental evidence for rG4 structure in the 3′UTR of *MAP3K10*

Circular dichroism (CD) is a spectroscopic method widely used to characterize rG4 structures (53). It measures the difference in the absorption of left-handed vs. right-handed circularly polarized light by a chiral molecule, such as a G4 DNA/RNA. We synthesized the predicted rG4 sequence (nucleotide positions 183-242 in the 3′UTR of *MAP3K10*, G4NN score ~ 0.97) by *in vitro* transcription. This RNA was subjected to CD spectroscopy. We observed a maximum ellipticity at ~264 nm and a minimum at ~240 nm, a characteristic feature of a parallel rG4 structure (54). Importantly, there was an increase (↑ 25.93 ± 4.8 %; N = 3 experiments) in the maximum value in the presence of potassium, which is known to stabilize rG4 structures. Another sequence of the same length derived from the initial part of the 3′UTR of *MAP3K10* (1 to 60 nucleotides, G4NN score 0.0042) did not show these rG4-specific spectral properties (Fig. 2A). TERRA rG4 sequence (10 repeats of UUAGGG), which served as positive control, also showed ellipticity maximum at ~264 nm and minimum at ~240 nm (Fig. 2B).

**Figure 2.**
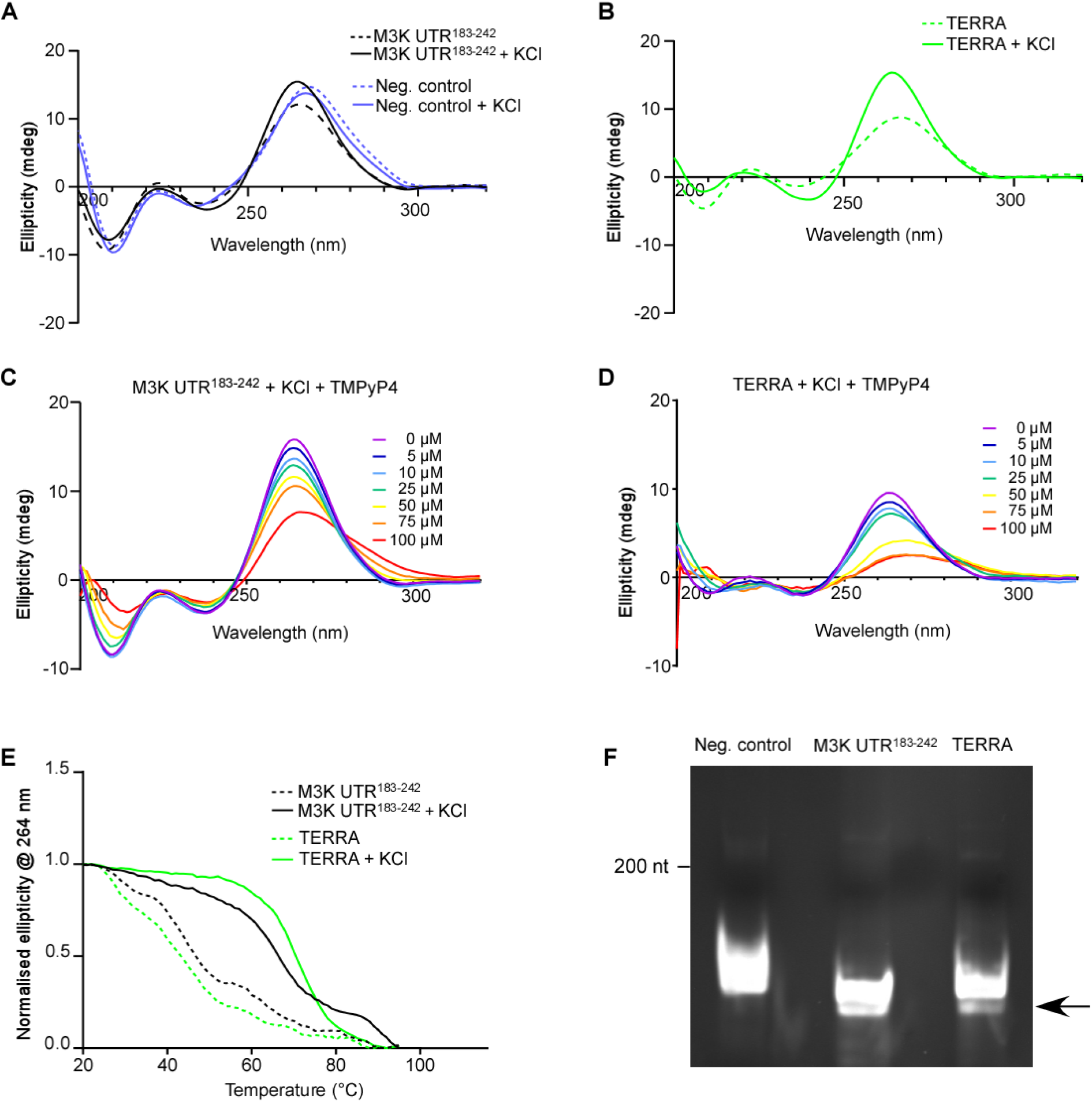
Experimental evidence for the rG4 structure in the 3′UTR of *MAP3K10*. (A) Circular Dichroism (CD) spectra of the RNA sequence derived from the predicted rG4 region of the 3′UTR of *MAP3K10* (183 to 242 nucleotides) with or without potassium chloride. (B) CD spectra of the TERRA sequence with or without potassium chloride. (C and D) CD spectra of the RNA sequence derived from the predicted rG4 region of the 3′UTR of *MAP3K10* (183 to 242 nucleotides) (C), and TERRA sequence (D), in the presence of different concentrations of TMPyP4 (Cayman). (E) CD values (λ 264nm) of indicated RNA sequences measured at different temperatures with or without potassium chloride (100 mM). (F) Native polyacrylamide gel electrophoresis showing the mobility of the indicated *in vitro* transcribed RNA sequences. Arrow indicates the fast-moving RNA species present in the test (3′UTR^183-242^ of *MAP3K10*) and positive control (TERRA). RNA sequence from 1 to 60 nucleotides of the 3′UTR of *MAP3K10* was used as a negative (neg.) control. Results are representatives of three (A, B, C, and F) or two (D and E) independent experiments.

The cationic porphyrin, 5,10,15,20-tetra(N-methyl-4-pyridyl)porphyrin (TMPyP4), is a G4 structure binding molecule. It stabilizes DNA G4s but unfolds RNA G4s (54,55). We investigated the effect of this molecule on rG4 of the 3′UTR of *MAP3K10* and the TERRA sequence. We observed a dose-dependent decrease in the maximum peak ellipticity values and an increase in the minimum peak ellipticity values, which suggests unfolding of the rG4 structure as previously reported (54) (Fig. 2C and D).

To assess the stability of the rG4 structure, we performed CD spectroscopy at temperatures ranging from 20°C to 95°C. We measured ellipticity at 264 nm (maxima) at these temperatures with or without potassium. The sequence derived from the 3′UTR of *MAP3K10* showed stable ellipticity till the physiological temperature (37°C) and beyond. In fact, its melting t_1/2_ was beyond 60°C in the presence of potassium. Importantly, the stability was reduced in the absence of potassium. The TERRA RNA was used as a positive control (Fig. 2E). These results provide evidence that the 3′UTR of *MAP3K10* can form an rG4 structure that is stable at physiological temperature in the presence of potassium.

Another method to investigate the presence of rG4 structure is the electrophoretic mobility shift assay. Intramolecular rG4 structures make the RNA (or DNA) sequences more compact and result in faster electrophoretic mobility under native conditions. We performed this assay on the predicted rG4 sequence (nucleotide position 183-242 in the 3′UTR of *MAP3K10*). The negative (1 to 60 nucleotides in the 3′UTR of *MAP3K10*) and positive (TERRA) controls described above were also used in this assay. In the positive control and the test sequence from *MAP3K10*, we observed a faster-moving band consistent with rG4 formation. This was not observed in the negative control (Fig. 2F). This observation provides another evidence for the ability of the 183-242 nucleotide region in the 3′UTR of *MAP3K10* to form an rG4 structure.

### The dORF in the 3′UTR of *MAP3K10* is translated

We next tested the translatability of the *MAP3K10* dORF experimentally using a luciferase-based reporter system. The coding sequence (CDS) of firefly luciferase (FLuc) was cloned downstream of the partial CDS of *MAP3K10* (717 nucleotides of the 3′ end) and its 3′UTR until the dORF (261 nucleotides after the canonical stop codon). The stop codon of the dORF and the start codon of the FLuc were removed so that translation initiation in the dORF, if it happens, will continue to generate firefly luciferase, whose activity can be quantified as luminescence (Fig. 3A). When transfected in HEK293 cells, this construct showed significant luciferase activity (2^nd^ bar in Fig. 3B). However, when the AUG of the dORF was mutated to UAC, the luciferase activity was reduced to the background level (3^rd^ bar in Fig. 3B). A construct without the 3′UTR served as a negative control. The levels of FLuc mRNA in transfected cells were comparable in all these conditions. These observations indicate that the dORF of *MAP3K10* has the potential to be translated.

**Figure 3.**
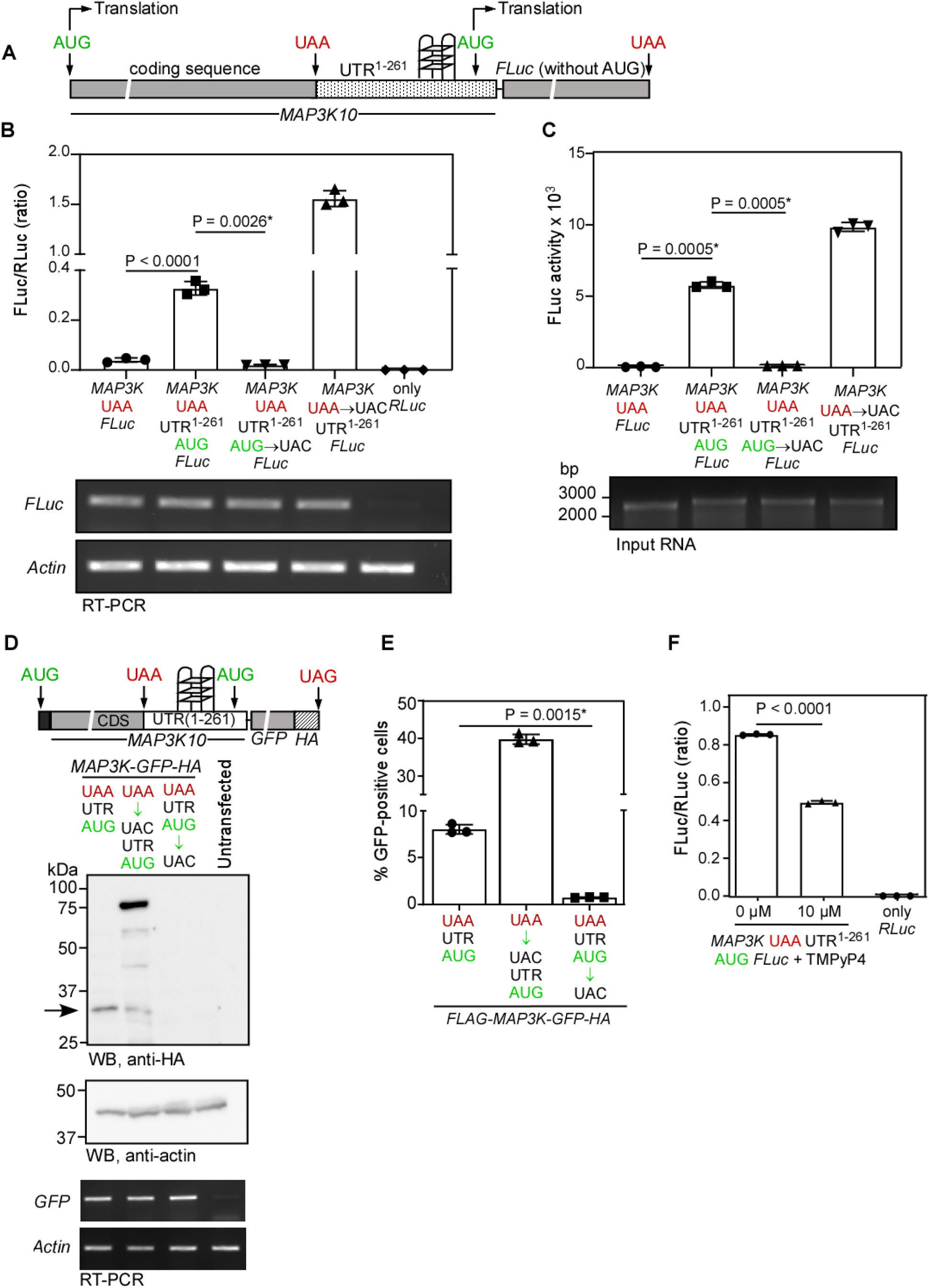
The dORF in the 3′UTR of *MAP3K10* is translated. (A) Schematic of the luciferase-based reporter used to investigate translation in the dORF. (B) Firefly luciferase (FLuc) activity (luminescence) in HEK293 cells transfected with the indicated constructs. FLuc activity was normalized with the activity of co-transfected *Renilla* luciferase, which was used as a transfection control. RT-PCR results indicating the FLuc mRNA levels are shown below. (C) FLuc activity obtained after *in vitro* transcription followed by *in vitro* translation of the indicated constructs. Rabbit reticulocyte lysate was used for *in vitro* translation. The input RNA is shown below. (D) Western blot image showing the product of dORF translation (arrow). HA-tagged GFP was used as a reporter in HEK293 cells (schematic above). Levels of GFP mRNA are shown below. The results are representative of three independent experiments. (E) Percentage of GFP-positive HEK293 cells transfected with the indicated reporter constructs and detected using a flow cytometer. (F) Normalized FLuc activity in HEK293 cells transfected with the indicated reporter construct and treated with TMPyP4 2h post-transfection. Bars in all graphs indicate mean ± SD (n=3 biological replicates). Statistical significance was calculated by a two-tailed unpaired Student’s t-test. *, Welch’s correction was applied.

To assess the translation efficiency of the dORF, we used another construct for comparison in which the FLuc was expressed via the canonical cap-dependent translation initiation from the 5′ end, in addition to the dORF translation. This was a variant of the construct shown in Figure 3A. The canonical stop codon of *MAP3K10* was mutated to UAC such that the canonical translation initiated at the 5′ end of the construct continues to generate FLuc. As expected, this construct showed much higher FLuc activity (4^th^ bar in Fig. 3B). The comparison revealed that the efficiency of the translation in the dORF was at least 21 ± 1.7 % of the canonical translation initiation at the 5′ end (2^nd^ vs 4^th^ bar in Fig. 3B). We observed similar results in an *in vitro* translation system based on rabbit reticulocyte lysate. Here, the efficiency of dORF translation was at least 58.7 ± 2.2 % of the canonical translation initiated at the 5′ end (Fig. 3C).

We next performed western blotting to detect the translation product from the dORF. For this, we used green fluorescent protein (GFP) as a reporter. The construct used was similar to the one described in Fig. 3A, except that the reporter used was HA-tagged GFP instead of FLuc (Schematic in Fig. 3D). When transfected in HEK293 cells, the construct yielded a HA-tagged protein of size between 37 and 25 kDa, which is consistent with the expected size (~31 kDa) of the product of translation initiated at the dORF (1^st^ lane Fig. 3D). The protein was absent when the AUG of the dORF was mutated to UAC (3^rd^ lane in Fig. 3D). Transfection of another construct where the canonical stop codon UAA was mutated to UAC yielded a larger protein expected from the translation initiated at the 5′ end. In addition, we also observed the protein generated by translation initiation in the dORF (2^nd^ lane in Fig. 3D). Furthermore, this assay ruled out stop codon readthrough (SCR) as the mechanism of translation of the dORF, as we did not detect the product of SCR (~ 68 kDa). The generation of GFP after the translation of the dORF was confirmed by quantifying the fluorescent cells using a flow cytometer (Fig. 3E). Altogether, these results obtained from luminescence-, fluorescence-, and western blotting-based reporter assays demonstrate that the dORF of *MAP3K10* is translated at significant levels (21% to 58%) relative to the canonical cap-mediated translation. We next investigated if this translation is driven by the rG4 structure present upstream of the dORF.

### The rG4 structure in the 3′UTR of *MAP3K10* drives the translation in the dORF

Evidence exists for 5′UTR rG4 structure–mediated cap-independent translation of *VEGFA, ARPC2*, and *FGF2* mRNAs (56–59). Since the rG4 structure in the 3′UTR of *MAP3K10* is upstream of the translatable dORF, we investigated whether this rG4 structure can drive the translation. First, we used TMPyP4, which melted the *MAP3K10* rG4 structure in our CD analysis (Fig. 2C). In the reporter assay described above (Fig. 3A), TMPyP4 reduced dORF translation, suggesting the importance of rG4 in this process (Fig. 3F).

We next employed a dual-luciferase assay to investigate the translation initiation potential of *MAP3K10* rG4. We cloned the *MAP3K10* rG4 sequence, followed by the dORF (181 to 261 nucleotides of the 3′UTR excluding the stop codon of the dORF), between the coding sequences of *Renilla* luciferase (RLuc) and firefly luciferase (FLuc). In this system, the translation of RLuc is canonical 5′ cap-mediated, whereas that of FLuc is driven by non-canonical mechanisms (termination-reinitiation, leaky scanning, and internal ribosome entry), if any, mediated by the sequence upstream of the FLuc (i.e., *MAP3K10* rG4 in our case) (Fig. 4A). The translation initiation ability of the *MAP3K10* rG4 sequence can be quantified using the FLuc to RLuc activity ratio.

**Figure 4.**
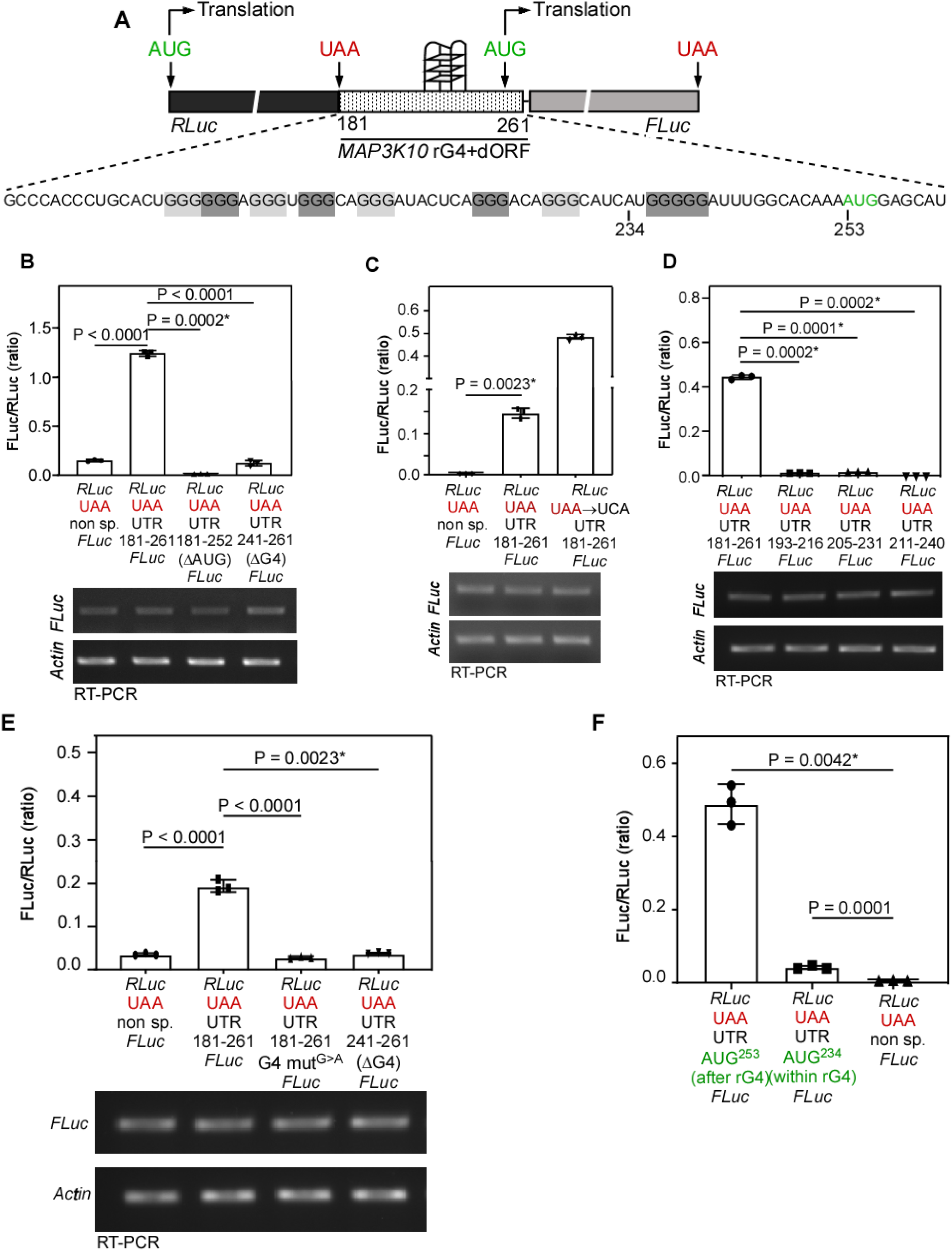
The rG4 structure in the 3′UTR of *MAP3K10* drives translation of the dORF. (A) Schematic of the dual luciferase-based reporter construct used to investigate the translation initiation activity of rG4. (B to F) Normalized FLuc activity (FLuc/RLuc) in HEK293 cells transfected with the indicated reporter constructs. Numbers next to UTR indicate the stretch of nucleotides in the 3′UTR cloned in the constructs. RT-PCR results indicating the FLuc mRNA levels are shown below the graphs. In deletion constructs used in (D), an AUG codon was introduced downstream of the deleted UTR in-frame with FLuc. Bars in all graphs indicate mean ± SD (n=3 biological replicates). Statistical significance was calculated by a two-tailed unpaired Student’s t-test. *, Welch’s correction was applied.

**Figure 5.**
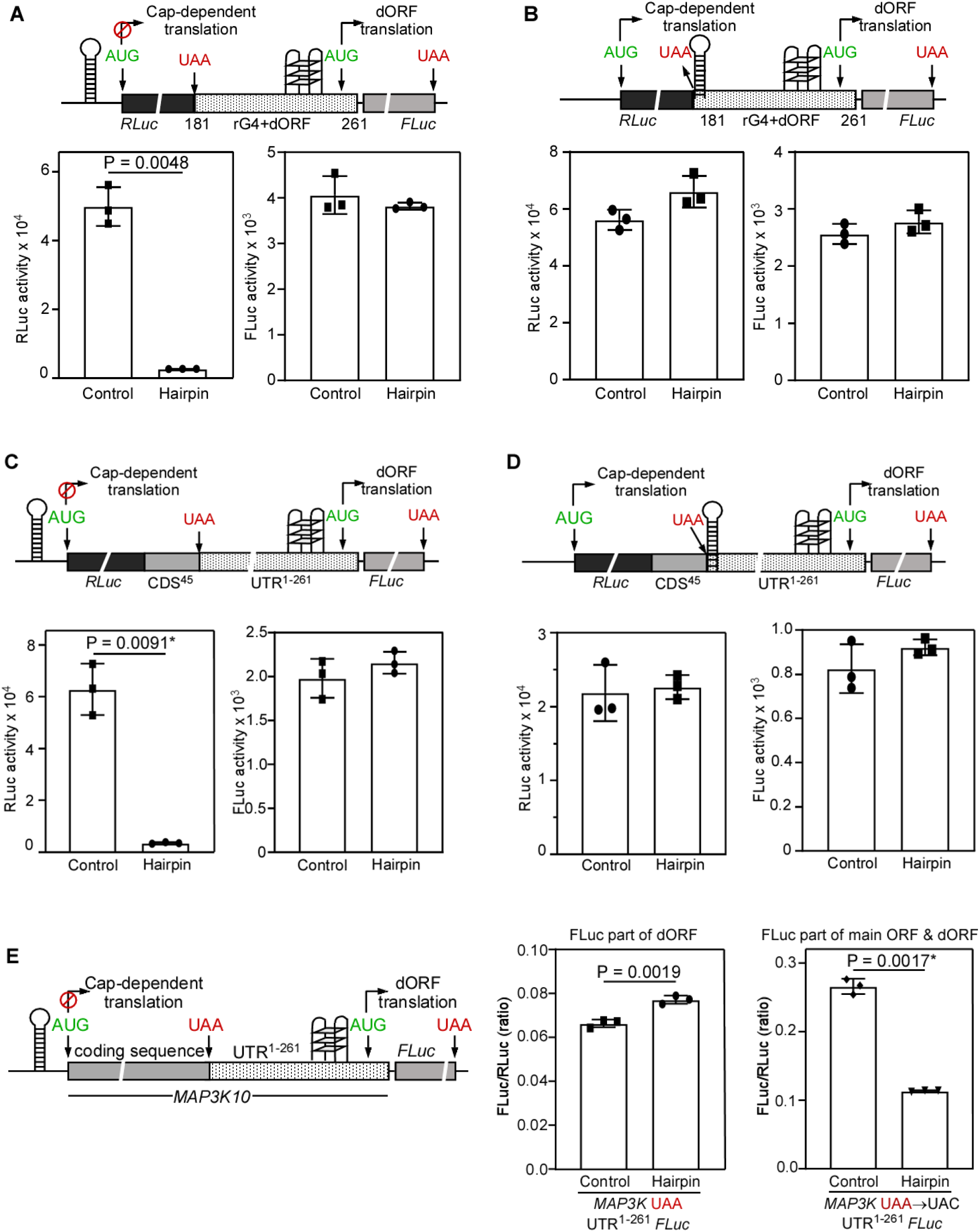
Translation of the dORF occurs independently of both the 5′ cap and the translation of the canonical ORF. (A to D) Activities of FLuc and Rluc in cells transfected with the indicated dual-luciferase constructs. (E) Activity of FLuc in cells transfected with the indicated single-luciferase construct. The FLuc activity was normalized to that of co-transfected RLuc. In (A), (C), and (E), a hairpin-forming sequence was placed upstream of the canonical ORF (RLuc or *MAP3K10*). In (B) and (D), the same hairpin-forming sequence was placed downstream of the canonical ORF (RLuc). Luciferase activities were measured 24 h after transfection. Bars in all graphs indicate mean ± SD (n=3 biological replicates). Statistical significance was calculated by a two-tailed unpaired Student’s t-test. *, Welch’s correction was applied.

When transfected in HEK293 cells, the *MAP3K10* rG4 sequence exhibited significant translation initiation activity, well above the background level observed with a negative control containing a non-specific sequence between RLuc and FLuc. The FLuc activity was reduced to background level when the AUG was removed, or the rG4 sequence was not included in the construct (Fig. 4B). RT-PCR analysis revealed comparable levels of FLuc mRNA in all the constructs used in the assay showing that the differences in the observed FLuc activities obtained from different constructs were not because of changes in the mRNA levels (Fig. 4B). RT-PCR performed using primers covering the 5′ end of RLuc and 3′ end of FLuc in cells transfected with the construct shown in Fig 4A resulted in a single expected amplicon of size 2870 bp. This observation provides evidence against cryptic alternative splicing of the mRNA that can potentially lead to FLuc expression without translation initiation at dORF (Fig. S2A).

The FLuc activity observed in these assays can also be due to cryptic promoter activity of the *MAP3K10* rG4. To test this, we performed a promoter activity assay using a promoter-less FLuc construct. When we cloned the *MAP3K10* rG4 sequence (1-261 nucleotides of the 3′UTR) into the upstream region of FLuc, we observed only background signal, indicating that the sequence lacks promoter activity. A construct without any promoter and another with the reverse sequence of rG4 served as negative controls to identify the background signal. The cytomegalovirus promoter was used as a positive control and showed robust FLuc activity (Fig. S2B). Furthermore, results from *in vitro* rabbit reticulocyte lysate-mediated translation assays provide additional evidence against cryptic promoter activity and cryptic splicing as explanations for dORF translation (Fig. 3C).

We next compared the efficiency of dORF translation by the *MAP3K10* rG4 sequence with that of 5′ cap-dependent canonical translation. For this, we used a variant of the construct shown in Fig. 4A, where the stop codon of RLuc UAA was mutated to UCA, such that the cap-mediated canonical translation leads to both RLuc and FLuc expression. When transfected in HEK293 cells, this construct showed robust FLuc activity, much above the FLuc activity obtained from the construct with the stop codon (UAA) of RLuc. The comparison of these two activities revealed that the translational initiation efficiency of the *MAP3K10* rG4 sequence was at least 29.4 ± 2.4 % of the canonical cap-mediated translation (Fig. 4C).

The *MAP3K10* rG4 sequence contains seven evolutionarily conserved stretches of Gs (≥ 3 Gs) (Fig. 1A). A minimum of 4 G-tracts is required to form a G4 structure. To test the minimum length of this sequence required for translation initiation activity, we generated three deletion variants (each had at least 4 G-tracts) of the construct depicted in Fig. 4A. When transfected in HEK293 cells, these constructs did not show any translation initiation ability, showing that all G-stretches are required for the translation initiation activity (Fig. 4D). Another construct where the 11 ‘G’s in the *MAP3K10* rG4 sequence were replaced by ‘A’s (G4 mut^G>A^), which disrupted all 7 G-tracts, also failed to show translation initiation activity (Fig. 4E). The AUG (A being 253^rd^ nucleotide in the 3′UTR of human *MAP3K10*) of the dORF is twelve nucleotides downstream of the last G-stretch. There is another AUG (A being the 234^th^ nucleotide), which overlaps with the last G-stretch (Fig. 4A). This is in +2 translation frame relative to that of dORF. We tested whether translation initiation can happen from this AUG (i.e., 234^th^ position). We made a variant of the construct shown in Fig. 4A where the FLuc was in frame with the AUG at the 234^th^ position. When transfected into HEK293 cells, we observed only background FLuc activity from the AUG at position 234. This shows that the distance between the rG4 sequence and the AUG is important for the translation initiation (Fig. 4F). Together, these results demonstrate that the *MAP3K10* rG4 sequence drives the translation of the dORF that encodes a tripeptide (Met-Glu-His). This is the second-shortest possible translation product, the first being just a dipeptide. Because of their short nature, it is nearly impossible to detect and annotate such peptides to a particular translatable ORF.

Some tripeptides have functional significance. For example, glutathione (Cys-Gly-Glu) is a potent intracellular antioxidant. This tripeptide is synthesized by non-ribosome-mediated enzymatic reactions (60). The tripeptide pyroglutamyl-histidyl-proline amide is the thyrotropin-releasing hormone synthesized and released from the hypothalamus to regulate the release of thyroid-stimulating hormone (TSH) from the anterior pituitary gland (61). The biological function, if any, of the tripeptide Met-Glu-His derived from the 3′UTR translon of *MAP3K10*, remains to be investigated.

### Translation of the dORF occurs independently of both the 5′ cap and the translation of the canonical ORF

We investigated whether the translation of the dORF is dependent on the 5′ cap-mediated translation of the canonical ORF. We used the dual luciferase assay described above (Fig. 4A). In this construct, a 42-nucleotide sequence previously shown to form a strong hairpin structure (ΔG = −41.1 kcal/mol) was inserted immediately upstream of RLuc. Another construct with an insert of the same length that does not form a hairpin structure was used as a negative control (schematic in Fig. 5A)(46). When transfected in cells, the construct with the hairpin yielded much lower RLuc expression than the one without it, consistent with inhibition of cap-mediated translation by the hairpin structure (46,62). However, there was no change in the rG4-mediated translation of the dORF harbouring FLuc (Fig. 5A).

We made another construct where the same hairpin structure was placed downstream of RLuc and upstream of the rG4 in the construct described above (Schematic in Fig. 5B). The hairpin structure can block the termination–reinitiation process as well as leaky scanning, if any. When transfected into cells, this construct did not show any change in FLuc expression, indicating that FLuc translation does not result from termination–reinitiation or leaky scanning (Fig. 5B).

Similar results were obtained when we performed these experiments in the presence of the partial coding sequence of *MAP3K10* (45 nucleotides) and its 3′UTR extending to the dORF (1-261 nucleotides). Presence of a hairpin upstream or downstream of the canonical ORF (i.e., RLuc) did not alter the translation of the dORF (i.e, FLuc) (Fig. 5C and D). Similar effects were observed when we replaced RLuc with the partial coding sequence of *MAP3K10*; the presence of the hairpin in its 5′UTR did not affect translation of the dORF, though it was able to affect the canonical translation (Fig. 5E). Also, the presence of the hairpin either upstream or downstream of the canonical ORF did not affect the levels of mRNA (Fig. S3A to D).

Collectively, these results show that the translation of the dORF of *MAP3K10* is independent of the 5′ cap and of the translation of the canonical ORF. Furthermore, since these hairpin structures can inhibit ribosomal scanning (62,63), these observations provide evidence against leaky ribosomal scanning as well as termination–reinitiation as mechanisms of translation of the dORF.

There are many examples of 5′ cap-independent translation in viruses. Here, translation is initiated by the internal entry of ribosomes mediated by RNA elements termed Internal Ribosome Entry Sites (IRES) (37). Although several such IRES elements are reported in mammalian cells (64–66), the extent of their activity is debated (40,67,68). We tested the potential IRES activity of *MAP3K10* rG4. We employed the Tornado (Twister-optimised RNA for durable overexpression) split nano-luciferase (nLuc) system for this (68). In this assay, the coding sequence of nLuc is split into a large bit (LgBiT) and a small bit (SmBiT) between which the sequence to be tested for IRES activity is cloned. Importantly, this split nLuc mRNA is flanked on both sides by ribozymes that mediate circularization of the mRNA following autocatalytic cleavage. If the test sequence has IRES activity, nLuc will be translated. In this assay, *MAP3K10* rG4 did not induce nLuc expression in HEK293 cells. However, known IRESs of encephalomyocarditis virus (EMCV) and Coxsackievirus B3 (CVB3) showed the expected activity and robust nLuc expression (Fig. S4). It is possible that the circularization of the mRNA disturbs the structure and function of the *MAP3K10* rG4. It should be noted that multiple reported cellular IRESs showed background activity in this assay in a previous study (68).

### Translation of the dORF regulates the expression of *MAP3K10*

The evolutionary conservation of the rG4 and its associated translatable dORF strongly suggests that this arrangement is functionally important in *MAP3K10*. Translation of ORFs present in UTRs can influence the translation of their corresponding canonical ORFs or their mRNA levels (8,33,69). To investigate this, we expressed FLAG-HA-tagged *MAP3K10* containing either the wild-type 3′UTR or a G4-mutant 3′UTR (G4 mut^G>A^, described above; 264 nucleotides) in HEK293 cells. We tested *MAP3K10* expression by western blotting and qRT-PCR. We observed a significant reduction in expression at both RNA and protein levels when expressed with a G4 mutant 3′UTR compared to that with the wild-type 3′UTR (Fig. 6A-C). These observations suggest a role of 3′UTR rG4-mediated translation in maintaining the levels of *MAP3K10*.

**Figure 6.**
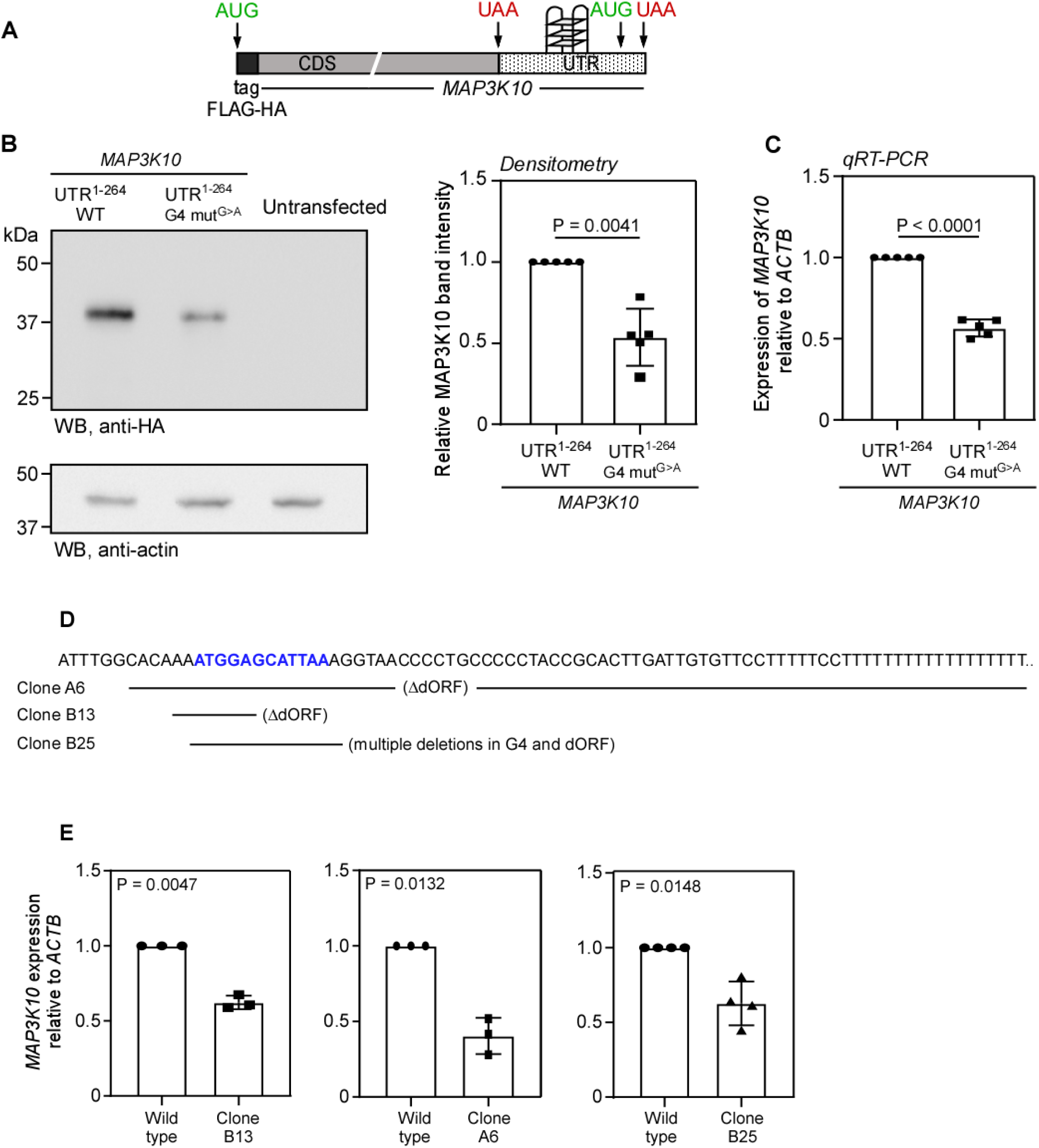
The rG4-driven translation of the dORF regulates the expression of *MAP3K10*. (A) Schematic of the construct expressing HA-tagged MAP3K10, including its 3′UTR. (B and C). The construct was transfected in HEK293 cells, and *MAP3K10* expression was detected by western blotting (B) and qRT-PCR (C) 24 h after transfection. (D) The sequence of the 3′UTR of *MAP3K10* around the dORF (blue). Regions deleted in multiple CRISPR clones are shown below the sequence, indicated by horizontal lines. (E) Results of qRT-PCR analysis showing the expression of *MAP3K10* in multiple CRISPR clones with deletions in the 3′UTR of *MAP3K10*. Bars in all graphs indicate mean ± SD (n=5 in B and C, n=3 or 4 in E, biological replicates). Statistical significance was calculated by a two-tailed paired Student’s t-test.

To investigate whether this regulation is functional in an endogenous context, we deleted the 3′UTR of endogenous *MAP3K10* using the CRISPR-Cas9 system in HEK293 cells. We obtained multiple heterozygous single-cell-derived clones with deletions in different stretches of the 3′UTR of *MAP3K10*. In these clones, the G4 region and/or the dORF were disrupted (Fig. S5). The clones where the dORF was disrupted (A6 and B13) and the clone where both the dORF and the G4 region were deleted (B25) showed reduced expression of *MAP3K10* (Fig. 6D-E). These results provide evidence that the translation in the dORF driven by an rG4 structure regulates *MAP3K10* expression.

*MAP3K10* (or *MLK2*) encodes a MAP kinase kinase kinase, a member of the serine/threonine kinase family. This kinase is activated by multiple forms of cellular stress, including UV irradiation and osmotic stress. Activated MAP3K10 phosphorylates mitogen-activated protein kinase 4 and 7 (MKK4 and MKK7), which in turn activate JNK (c-Jun N-terminal kinases), leading to apoptosis or cell survival depending on the context and the extent of activation (70–72). The expression of *MAP3K10* varies across different human tissues. According to the Protein Atlas (proteinatlas.org), the MAP3K10 protein is highly expressed in skeletal muscles, lymphoid tissue, the kidney, and the gastrointestinal tract. In the brain, *MAP3K10* mRNA levels are highest, although the protein levels are moderate compared to other tissues. Our experimental results show that this translon positively regulates the levels of *MAP3K10* mRNA. Together, these observations suggest that the 3′UTR translon of *MAP3K10* might play a critical role in tissue-specific expression of *MAP3K10*.

Previous observations indicated that translation of ORFs in the 3′UTR can have functions independent of the polypeptides generated from translation (8,33). Our observations support this idea. HEK293 cells that lack the dORF or the rG4 or both in the 3′UTR of *MAP3K10* show reduced levels of *MAP3K10* mRNA. The exact mechanism by which this is caused remains to be investigated. It is possible that the translating ribosomes protect the mRNA from degradation by nucleases or miRNAs. In fact, the predicted binding site of hsa-miR-155-5p coincides with the 3′UTR translon of *MAP3K10* (73,74). In the absence of translation in the 3′UTR, the mRNA may become susceptible to degradation by these factors. This hypothesis of the mRNA protection function of translation in the 3′UTR remains to be tested.

We believe that this study provides important insights into the ongoing debate surrounding mechanisms of internal translation initiation in eukaryotic mRNAs. Several plausible mechanisms could account for translation of the MAP3K10 dORF, and we have systematically evaluated each of them. Western blotting and 3′ hairpin assays effectively rule out stop-codon readthrough as a source of dORF expression. Moreover, the combined results of the 5′ and 3′ hairpin experiments do not support mechanisms such as leaky scanning, ribosome shunting, or termination-reinitiation. Potential artifacts, including cryptic promoter activity and cryptic splicing, were excluded through in vitro translation assays, RT–PCR analyses, and promoterless reporter controls. Collectively, these findings are most consistent with an internal mode of translation initiation that is independent of the 5′ cap.

## ACKNOWLEDGEMENTS

This work was supported by funding from the Swarnajayanti Fellowship (DST/SJF/LSA-04/2019-20) awarded by the Department of Science and Technology (DST). Additional support was provided by the STARS grant from the Ministry of Education (STARS/APR2019/BS/328/FS), DBT’s Genome Engineering Technology Program (BT/PR38405/GET/119/309/2020), the Indian Council of Medical Research, Anusandhan National Research Foundation, the Blockchain Foundation of India, the EMBO Global Investigator Network, DST Funds for Improvement of Science and Technology infrastructure, and the Institute of Eminence funds allocated by the Ministry of Education to the Indian Institute of Science.

## AUTHOR CONTRIBUTIONS

**Kirtana Vasu:** Conceptualization; Data curation; Formal analysis; Investigation; Methodology. **Swati Ghosh, Sarthak Sahoo:** Investigation; Methodology. **Debraj Manna, Saubhik Som, Md Noor Akhtar and Debaleena Kar:** Resources; Methodology. **Sandeep M Eswarappa:** Conceptualization; Resources; Data curation; Formal analysis; Supervision; Funding acquisition; Investigation; Methodology; Writing—original draft; Project administration.

## Data Availability

All the data related to this study are provided with the manuscript

## REFERENCES

1. Slavoff, S.A., Mitchell, A.J., Schwaid, A.G., Cabili, M.N., Ma, J., Levin, J.Z., Karger, A.D., Budnik, B.A., Rinn, J.L. and Saghatelian, A. (2013) Peptidomic discovery of short open reading frame-encoded peptides in human cells. Nat Chem Biol, 9, 59–64.

2. Chen, J., Brunner, A.D., Cogan, J.Z., Nunez, J.K., Fields, A.P., Adamson, B., Itzhak, D.N., Li, J.Y., Mann, M., Leonetti, M.D. et al. (2020) Pervasive functional translation of noncanonical human open reading frames. Science, 367, 1140–1146.

3. Jurgens, L. and Wethmar, K. (2022) The Emerging Role of uORF-Encoded uPeptides and HLA uLigands in Cellular and Tumor Biology. Cancers (Basel), 14.

4. Ingolia, N.T., Brar, G.A., Stern-Ginossar, N., Harris, M.S., Talhouarne, G.J., Jackson, S.E., Wills, M.R. and Weissman, J.S. (2014) Ribosome profiling reveals pervasive translation outside of annotated protein-coding genes. Cell Rep, 8, 1365–1379.

5. Ingolia, N.T., Lareau, L.F. and Weissman, J.S. (2011) Ribosome profiling of mouse embryonic stem cells reveals the complexity and dynamics of mammalian proteomes. Cell, 147, 789–802.

6. Crappe, J., Van Criekinge, W., Trooskens, G., Hayakawa, E., Luyten, W., Baggerman, G. and Menschaert, G. (2013) Combining in silico prediction and ribosome profiling in a genome-wide search for novel putatively coding sORFs. BMC Genomics, 14, 648.

7. Bazzini, A.A., Johnstone, T.G., Christiano, R., Mackowiak, S.D., Obermayer, B., Fleming, E.S., Vejnar, C.E., Lee, M.T., Rajewsky, N., Walther, T.C. et al. (2014) Identification of small ORFs in vertebrates using ribosome footprinting and evolutionary conservation. EMBO J, 33, 981–993.

8. Wu, Q., Wright, M., Gogol, M.M., Bradford, W.D., Zhang, N. and Bazzini, A.A. (2020) Translation of small downstream ORFs enhances translation of canonical main open reading frames. EMBO J, 39, e104763.

9. Swirski, M.I., Tierney, J.A.S., Alba, M.M., Andreev, D.E., Aspden, J.L., Atkins, J.F., Bassani-Sternberg, M., Berry, M.J., Biffo, S., Boris-Lawrie, K. et al. (2025) Translon: a single term for translated regions. Nat Methods.

10. Johnstone, T.G., Bazzini, A.A. and Giraldez, A.J. (2016) Upstream ORFs are prevalent translational repressors in vertebrates. EMBO J, 35, 706–723.

11. Calvo, S.E., Pagliarini, D.J. and Mootha, V.K. (2009) Upstream open reading frames cause widespread reduction of protein expression and are polymorphic among humans. Proc Natl Acad Sci U S A, 106, 7507–7512.

12. Ye, Y., Liang, Y., Yu, Q., Hu, L., Li, H., Zhang, Z. and Xu, X. (2015) Analysis of human upstream open reading frames and impact on gene expression. Hum Genet, 134, 605–612.

13. Calkhoven, C.F., Muller, C. and Leutz, A. (2000) Translational control of C/EBPalpha and C/EBPbeta isoform expression. Genes Dev, 14, 1920–1932.

14. Kochetov, A.V., Ahmad, S., Ivanisenko, V., Volkova, O.A., Kolchanov, N.A. and Sarai, A. (2008) uORFs, reinitiation and alternative translation start sites in human mRNAs. FEBS Lett, 582, 1293–1297.

15. Bastide, A., Karaa, Z., Bornes, S., Hieblot, C., Lacazette, E., Prats, H. and Touriol, C. (2008) An upstream open reading frame within an IRES controls expression of a specific VEGF-A isoform. Nucleic Acids Res, 36, 2434–2445.

16. Mueller, P.P. and Hinnebusch, A.G. (1986) Multiple upstream AUG codons mediate translational control of GCN4. Cell, 45, 201–207.

17. Ebina, I., Takemoto-Tsutsumi, M., Watanabe, S., Koyama, H., Endo, Y., Kimata, K., Igarashi, T., Murakami, K., Kudo, R., Ohsumi, A. et al. (2015) Identification of novel Arabidopsis thaliana upstream open reading frames that control expression of the main coding sequences in a peptide sequence-dependent manner. Nucleic Acids Res, 43, 1562–1576.

18. Rahmani, F., Hummel, M., Schuurmans, J., Wiese-Klinkenberg, A., Smeekens, S. and Hanson, J. (2009) Sucrose control of translation mediated by an upstream open reading frame-encoded peptide. Plant Physiol, 150, 1356–1367.

19. Starck, S.R., Tsai, J.C., Chen, K., Shodiya, M., Wang, L., Yahiro, K., Martins-Green, M., Shastri, N. and Walter, P. (2016) Translation from the 5’ untranslated region shapes the integrated stress response. Science, 351, aad3867.

20. Cloutier, P., Poitras, C., Faubert, D., Bouchard, A., Blanchette, M., Gauthier, M.S. and Coulombe, B. (2020) Upstream ORF-Encoded ASDURF Is a Novel Prefoldin-like Subunit of the PAQosome. J Proteome Res, 19, 18–27.

21. Lawless, C., Pearson, R.D., Selley, J.N., Smirnova, J.B., Grant, C.M., Ashe, M.P., Pavitt, G.D. and Hubbard, S.J. (2009) Upstream sequence elements direct post-transcriptional regulation of gene expression under stress conditions in yeast. BMC Genomics, 10, 7.

22. Janich, P., Arpat, A.B., Castelo-Szekely, V., Lopes, M. and Gatfield, D. (2015) Ribosome profiling reveals the rhythmic liver translatome and circadian clock regulation by upstream open reading frames. Genome Res, 25, 1848–1859.

23. Wethmar, K., Schulz, J., Muro, E.M., Talyan, S., Andrade-Navarro, M.A. and Leutz, A. (2016) Comprehensive translational control of tyrosine kinase expression by upstream open reading frames. Oncogene, 35, 1736–1742.

24. Dasgupta, A. and Prensner, J.R. (2024) Upstream open reading frames: new players in the landscape of cancer gene regulation. NAR Cancer, 6, zcae023.

25. Sun, Y., Shui, K., Li, Q., Liu, C., Jin, W., Ni, J.Q., Lu, J. and Zhang, L. (2025) Upstream open reading frames dynamically modulate CLOCK protein translation to regulate circadian rhythms and sleep. PLoS Biol, 23, e3003173.

26. Lee, D.S.M., Park, J., Kromer, A., Baras, A., Rader, D.J., Ritchie, M.D., Ghanem, L.R. and Barash, Y. (2021) Disrupting upstream translation in mRNAs is associated with human disease. Nat Commun, 12, 1515.

27. Silva, J., Fernandes, R. and Romao, L. (2019) Translational Regulation by Upstream Open Reading Frames and Human Diseases. Adv Exp Med Biol, 1157, 99–116.

28. Yang, H., Li, Q., Stroup, E.K., Wang, S. and Ji, Z. (2024) Widespread stable noncanonical peptides identified by integrated analyses of ribosome profiling and ORF features. Nat Commun, 15, 1932.

29. Hsu, P.Y., Calviello, L., Wu, H.L., Li, F.W., Rothfels, C.J., Ohler, U. and Benfey, P.N. (2016) Super-resolution ribosome profiling reveals unannotated translation events in Arabidopsis. Proc Natl Acad Sci U S A, 113, E7126–E7135.

30. Chothani, S.P., Adami, E., Widjaja, A.A., Langley, S.R., Viswanathan, S., Pua, C.J., Zhihao, N.T., Harmston, N., D’Agostino, G., Whiffin, N. et al. (2022) A high-resolution map of human RNA translation. Mol Cell, 82, 2885–2899 e2888.

31. Hu, F., Lu, J., Matheson, L.S., Diaz-Munoz, M.D., Saveliev, A., Xu, J. and Turner, M. (2021) ORFLine: a bioinformatic pipeline to prioritize small open reading frames identifies candidate secreted small proteins from lymphocytes. Bioinformatics, 37, 3152–3159.

32. Ji, Z., Song, R., Regev, A. and Struhl, K. (2015) Many lncRNAs, 5’UTRs, and pseudogenes are translated and some are likely to express functional proteins. Elife, 4, e08890.

33. Nobuta, R., Machida, K., Sato, M., Hashimoto, S., Toriumi, Y., Nakajima, S., Suto, D., Imataka, H. and Inada, T. (2020) eIF4G-driven translation initiation of downstream ORFs in mammalian cells. Nucleic Acids Res, 48, 10441–10455.

34. Skabkin, M.A., Skabkina, O.V., Hellen, C.U. and Pestova, T.V. (2013) Reinitiation and other unconventional posttermination events during eukaryotic translation. Mol Cell, 51, 249–264.

35. Hinnebusch, A.G. (1997) Translational regulation of yeast GCN4. A window on factors that control initiator-trna binding to the ribosome. J Biol Chem, 272, 21661–21664.

36. Vattem, K.M. and Wek, R.C. (2004) Reinitiation involving upstream ORFs regulates ATF4 mRNA translation in mammalian cells. Proc Natl Acad Sci U S A, 101, 11269–11274.

37. Jaafar, Z.A. and Kieft, J.S. (2019) Viral RNA structure-based strategies to manipulate translation. Nat Rev Microbiol, 17, 110–123.

38. Yang, Y. and Wang, Z. (2019) IRES-mediated cap-independent translation, a path leading to hidden proteome. J Mol Cell Biol, 11, 911–919.

39. Lacerda, R., Menezes, J. and Romao, L. (2017) More than just scanning: the importance of cap-independent mRNA translation initiation for cellular stress response and cancer. Cell Mol Life Sci, 74, 1659–1680.

40. Koch, P., Zhang, Z., Genuth, N.R., Susanto, T.T., Haimann, M., Khmelinskaia, A., Byeon, G.W., Dey, S., Barna, M. and Leppek, K. (2025) A versatile toolbox for determining IRES activity in cells and embryonic tissues. EMBO J, 44, 2695–2724.

41. Fay, M.M., Lyons, S.M. and Ivanov, P. (2017) RNA G-Quadruplexes in Biology: Principles and Molecular Mechanisms. J Mol Biol, 429, 2127–2147.

42. Lyu, K., Chow, E.Y., Mou, X., Chan, T.F. and Kwok, C.K. (2021) RNA G-quadruplexes (rG4s): genomics and biological functions. Nucleic Acids Res, 49, 5426–5450.

43. Dhapola, P. and Chowdhury, S. (2016) QuadBase2: web server for multiplexed guanine quadruplex mining and visualization. Nucleic Acids Res, 44, W277–283.

44. Garant, J.M., Perreault, J.P. and Scott, M.S. (2018) G4RNA screener web server: User focused interface for RNA G-quadruplex prediction. Biochimie, 151, 115–118.

45. Eswarappa, S.M., Potdar, A.A., Koch, W.J., Fan, Y., Vasu, K., Lindner, D., Willard, B., Graham, L.M., DiCorleto, P.E. and Fox, P.L. (2014) Programmed translational readthrough generates antiangiogenic VEGF-Ax. Cell, 157, 1605–1618.

46. Babendure, J.R., Babendure, J.L., Ding, J.H. and Tsien, R.Y. (2006) Control of mammalian translation by mRNA structure near caps. RNA, 12, 851–861.

47. Manjunath, L.E., Singh, A., Som, S. and Eswarappa, S.M. (2023) Mammalian proteome expansion by stop codon readthrough. Wiley Interdiscip Rev RNA, 14, e1739.

48. Garant, J.M., Perreault, J.P. and Scott, M.S. (2017) Motif independent identification of potential RNA G-quadruplexes by G4RNA screener. Bioinformatics, 33, 3532–3537.

49. Gruber, A.R., Lorenz, R., Bernhart, S.H., Neubock, R. and Hofacker, I.L. (2008) The Vienna RNA websuite. Nucleic Acids Res, 36, W70–74.

50. Biffi, G., Tannahill, D. and Balasubramanian, S. (2012) An intramolecular G-quadruplex structure is required for binding of telomeric repeat-containing RNA to the telomeric protein TRF2. J Am Chem Soc, 134, 11974–11976.

51. Huppert, J.L., Bugaut, A., Kumari, S. and Balasubramanian, S. (2008) G-quadruplexes: the beginning and end of UTRs. Nucleic Acids Res, 36, 6260–6268.

52. Kwok, C.K., Marsico, G., Sahakyan, A.B., Chambers, V.S. and Balasubramanian, S. (2016) rG4-seq reveals widespread formation of G-quadruplex structures in the human transcriptome. Nat Methods, 13, 841–844.

53. Del Villar-Guerra, R., Trent, J.O. and Chaires, J.B. (2018) G-Quadruplex Secondary Structure Obtained from Circular Dichroism Spectroscopy. Angew Chem Int Ed Engl, 57, 7171–7175.

54. Morris, M.J., Wingate, K.L., Silwal, J., Leeper, T.C. and Basu, S. (2012) The porphyrin TmPyP4 unfolds the extremely stable G-quadruplex in MT3-MMP mRNA and alleviates its repressive effect to enhance translation in eukaryotic cells. Nucleic Acids Res, 40, 4137–4145.

55. Zamiri, B., Reddy, K., Macgregor, R.B., Jr. and Pearson, C.E. (2014) TMPyP4 porphyrin distorts RNA G-quadruplex structures of the disease-associated r(GGGGCC)n repeat of the C9orf72 gene and blocks interaction of RNA-binding proteins. J Biol Chem, 289, 4653–4659.

56. Morris, M.J., Negishi, Y., Pazsint, C., Schonhoft, J.D. and Basu, S. (2010) An RNA G-quadruplex is essential for cap-independent translation initiation in human VEGF IRES. J Am Chem Soc, 132, 17831–17839.

57. Bhattacharyya, D., Diamond, P. and Basu, S. (2015) An Independently folding RNA G-quadruplex domain directly recruits the 40S ribosomal subunit. Biochemistry, 54, 1879–1885.

58. Bonnal, S., Schaeffer, C., Creancier, L., Clamens, S., Moine, H., Prats, A.C. and Vagner, S. (2003) A single internal ribosome entry site containing a G quartet RNA structure drives fibroblast growth factor 2 gene expression at four alternative translation initiation codons. J Biol Chem, 278, 39330–39336.

59. Al-Zeer, M.A., Dutkiewicz, M., von Hacht, A., Kreuzmann, D., Rohrs, V. and Kurreck, J. (2019) Alternatively spliced variants of the 5’-UTR of the ARPC2 mRNA regulate translation by an internal ribosome entry site (IRES) harboring a guanine-quadruplex motif. RNA Biol, 16, 1622–1632.

60. Meister, A. (1988) Glutathione metabolism and its selective modification. J Biol Chem, 263, 17205–17208.

61. Fliers, E., Wiersinga, W.M. and Swaab, D.F. (1998) Physiological and pathophysiological aspects of thyrotropin-releasing hormone gene expression in the human hypothalamus. Thyroid, 8, 921–928.

62. Kozak, M. (1986) Influences of mRNA secondary structure on initiation by eukaryotic ribosomes. Proc Natl Acad Sci U S A, 83, 2850–2854.

63. Kozak, M. (1989) Circumstances and mechanisms of inhibition of translation by secondary structure in eucaryotic mRNAs. Mol Cell Biol, 9, 5134–5142.

64. Weingarten-Gabbay, S., Elias-Kirma, S., Nir, R., Gritsenko, A.A., Stern-Ginossar, N., Yakhini, Z., Weinberger, A. and Segal, E. (2016) Comparative genetics. Systematic discovery of cap-independent translation sequences in human and viral genomes. Science, 351.

65. Komar, A.A. and Hatzoglou, M. (2011) Cellular IRES-mediated translation: the war of ITAFs in pathophysiological states. Cell Cycle, 10, 229–240.

66. Xue, S., Tian, S., Fujii, K., Kladwang, W., Das, R. and Barna, M. (2015) RNA regulons in Hox 5’ UTRs confer ribosome specificity to gene regulation. Nature, 517, 33–38.

67. Akirtava, C., May, G.E. and McManus, C.J. (2022) False-positive IRESes from Hoxa9 and other genes resulting from errors in mammalian 5’ UTR annotations. Proc Natl Acad Sci U S A, 119, e2122170119.

68. Unti, M.J., Doetsch, L. and Jaffrey, S.R. (2024) A circular split nanoluciferase reporter for validating and screening putative internal ribosomal entry site elements. RNA, 30, 1529–1540.

69. Musaev, D., Abdelmessih, M., Vejnar, C.E., Yartseva, V., Weiss, L.A., Strayer, E.C., Takacs, C.M. and Giraldez, A.J. (2024) UPF1 regulates mRNA stability by sensing poorly translated coding sequences. Cell Rep, 43, 114074.

70. Hirai, S., Katoh, M., Terada, M., Kyriakis, J.M., Zon, L.I., Rana, A., Avruch, J. and Ohno, S. (1997) MST/MLK2, a member of the mixed lineage kinase family, directly phosphorylates and activates SEK1, an activator of c-Jun N-terminal kinase/stress-activated protein kinase. J Biol Chem, 272, 15167–15173.

71. Mordente, K., Ryder, L. and Bekker-Jensen, S. (2024) Mechanisms underlying sensing of cellular stress signals by mammalian MAP3 kinases. Mol Cell, 84, 142–155.

72. Gallo, K.A. and Johnson, G.L. (2002) Mixed-lineage kinase control of JNK and p38 MAPK pathways. Nat Rev Mol Cell Biol, 3, 663–672.

73. Wang, C., Zhang, X., Zhang, C., Zhai, F., Li, Y. and Huang, Z. (2017) MicroRNA-155 targets MAP3K10 and regulates osteosarcoma cell growth. Pathol Res Pract, 213, 389–393.

74. Zhu, J., Chen, T., Yang, L., Li, Z., Wong, M.M., Zheng, X., Pan, X., Zhang, L. and Yan, H. (2012) Regulation of microRNA-155 in atherosclerotic inflammatory responses by targeting MAP3K10. PLoS One, 7, e46551.

